# Root-specific activation of plasma membrane H^+^-ATPase 1 enhances plant growth and shoot accumulation of nutrient elements under nutrient-poor conditions in *Arabidopsis thaliana*

**DOI:** 10.1101/2022.05.19.492745

**Authors:** Kota Monden, Takehiro Kamiya, Daisuke Sugiura, Takamasa Suzuki, Tsuyoshi Nakagawa, Takushi Hachiya

## Abstract

Plasma membrane (PM) H^+^-ATPase contributes to nutrient uptake and stomatal opening by creating proton gradient across the membrane. Previous studies report that a dominant mutation in the *OPEN STOMATA2* locus (*OST2-2D*) constitutively activates *Arabidopsis* PM H^+^-ATPase 1 (AHA1), which enlarges proton motive force for root nutrient uptake. However, the stomatal opening is also constitutively enhanced in the *ost2-2D*, causing drought hypersensitivity. To develop plants with improved nutrient acquisition and normal stomatal movement, we generated grafted plants (scion/rootstock : Col-0 (WT)/*ost2-2D*), and compared their growth and nutrient element content with those of control plants (WT/WT) under two nutrient regimes. WT/*ost2-2D* shoots had larger weights, rosette diameter, leaf blade area, and content of C, N, K, Ca, S, P, Mg, Na, Mn, B, Co, and Mo under nutrient-poor conditions compared with WT/WT shoots. The root weights and primary root length were greater in WT/*ost2-2D* plants than in WT/WT plants under both nutrient conditions. These results suggest that root-specific activation of PM H^+^-ATPase enhances root nutrient uptake, which accelerates the plant growth under nutrient-poor condition. Our study presents a novel approach to improving nutrient uptake efficiency in crops and vegetable for the implementation of low-input sustainable agriculture.

## 1. Introduction

A growing population requires agriculture to increase crop production on a global scale. Nutrient availability is an important determinant of crop productivity; thus, fertilizers are often applied in large amounts to improve crop production. Intensive fertilizer use, however, puts a heavy burden on the environment, e.g., through groundwater pollution, eutrophication, and greenhouse gas emissions [1]. N, P, and K, macronutrients typically in high demand in plants, account for a major portion of applied fertilizers. The artificial synthesis of N fertilizers requires massive fossil fuel consumption; i.e., ammonia synthesis via the Harber–Bosch process consumes 1%–2% of the global energy budget currently [2]. Additionally, the P contained in fertilizers is derived from nonrenewable rock phosphate that will be depleted in the future [3]. Thus, developing plants with high nutrient uptake efficiency (NUE) is crucial to achieving sustainable agriculture.

It is widely accepted that plasma membrane (PM) H^+^-ATPase provides proton motive force for root nutrient uptake mediated by specific transporters and channels [4]. Constitutive overexpression of *Oryza sativa* PM H^+^-ATPase 1 (*OSA1*) under the control of the *CaMV-35S* promoter was shown to enhance root uptake of ammonium, plant growth, and productivity in both the field and laboratory [5]. Conversely, similar overexpression of PM H^+^-ATPase genes failed to confer any positive effects on *Arabidopsis* and tobacco plants [6, 7]. Importantly, post-translational activation of PM H^+^-ATPase often has major effects on *Arabidopsis* physiological responses, including stomatal opening and hypocotyl elongation [8, 9]. Therefore, with the aim of improving the NUE of *Arabidopsis*, we focused on the *Arabidopsis OPEN STOMATA 2* (*OST2*)-dominant mutant (*ost2-2D*), which expresses a constitutively active allele of the *Arabidopsis* PM H^+^-ATPase 1 (*AHA1*) [9]. This mutant exhibits lower apoplastic pH and longer-sized root epidermal cells, corresponding to enhancements in proton motive force and acid growth in the roots, respectively [10]. However, PM hyperpolarization in the guard cells of *ost2-2D* causes drought hypersensitivity due to the prevention of stomatal closure, even with the addition of abscisic acid [9, 11, 12]. Thus, the root-specific activation of OST2/AHA1 could be an ideal design for improving NUE. To achieve this design, we grafted Col-0 (wild-type; WT) scions onto rootstock originating from WT or *ost2-2D* and analyzed the vegetative growth, nutrient element content, and transcriptomes of the grafted plants grown under nutrient-rich or nutrient-poor conditions. We show that *ost2-2D*-derived rootstock enhances the growth of shoots and roots as well as the shoot accumulation of several nutrient elements under nutrient-poor conditions.

## 2. Materials and methods

### 2.1. Plant materials and growth conditions

Seeds of Col-0 (WT) and *ost2-2D* [9] were surface sterilized and sown in a solid medium containing 50 mL of half-strength Murashige and Skoog (1/2-MS) salts including vitamins (Code M0222; Duchefa Biochemie, Haarlem, The Netherlands) supplemented with 0.05% (w/v) MES-KOH (pH 5.7), 1% (w/v) sucrose, and 0.5% (w/v) gellan gum (Fujifilm Wako, Osaka, Japan). The seeds were placed in the dark at 4°C for 3 days to break dormancy. Seedlings were grown vertically under a PPFD of 25 μmol m^-2^ s^-1^ (16/8-h light/dark cycle) at 22°C. Micrografting was performed aseptically using 5-day-old seedlings [13], which were cut perpendicularly in the hypocotyl with an injection needle tip (Code NN-2613S; TERUMO, Tokyo, Japan). The obtained scion was connected with the partner rootstock in a silicon microtube (Code 1-8194-04; AS ONE, Osaka, Japan). The grafted plants were incubated for 5 days under a PPDF of 50 μmol m^-2^ s^-1^ (constant light) at 27°C and further cultivated for 1 day under a PPFD of 25 μmol m^-2^ s^-1^ (16/8-h light/dark cycle) at 22°C except that the grafted plants used for the measurement of root length were not subjected to cultivation for 1 day under a PPFD of 25 μmol m^-2^ s^-1^ (16/8-h light/dark cycle) at 22°C. Successfully grafted plants without adventitious root formation were selected for similar size; transferred to vermiculite pots, soil pots, or solid media; and grown for use in the following experiments. For vermiculite cultivation, the grafted plants were grown under a PPFD of 100–120 μmol m^-2^ s^-1^ (16/8-h light/dark cycle) at 22°C–23°C. Before the cultivation, ~150 mL of vermiculite was soaked with 80 mL of 1/2-MS or 1/50-MS salts without vitamins (Code M0221; Duchefa Biochemie) supplemented with 0.05% (w/v) MES-KOH (pH 5.7). Four days after the onset of cultivation, 30 mL of the same compositions of 1/2-MS or 1/50-MS salts was supplied to the vermiculite. The plants were then cultivated for an additional 3 days; after which, they were sampled for experiments. For soil cultivation, the grafted plants were grown for 7 days under a PPFD of 100–120 μmol m^-2^ s^-1^ (16/8-h light/dark cycle) at 22°C–23°C in the pots containing ~150 mL of nutrient-rich soil (Supermix A; Sakata Seed, Kyoto, Japan). For an *in vitro* culture on solid media, the plants were grown for 3 days (vertically) or 7 days (horizontally) under a PPFD of 100–120 μmol m^-2^ s^-1^ (16/8-h light/dark cycle) at 22°C–23°C in 30 mL (horizontally) or 50 mL (vertically) of 1/2-MS or 1/50-MS salts without vitamins (Duchefa Biochemie) supplemented with 0.05% (w/v) MES-KOH (pH 5.7), 1% (w/v) sucrose, and 0.8% (w/v) purified agar (Code 01162–15; Nacalai Tesque, Kyoto, Japan). To allow nondestructive sampling of the roots, the surfaces of solid media were covered with cellophane sheets [14]. All other details are described in the figure legends.

### 2.2. Growth analysis of plants under vermiculite and soil cultivation

Shoots were weighed on a precision balance (HR-202i; A&D, Tokyo, Japan) and scanned at a resolution of 600 dpi (GT-X980; EPSON, Tokyo, Japan) to measure rosette diameter. Leaf blades were harvested from the shoots and scanned at a resolution of 600 dpi to measure the area of the leaf blades. The shoots were then oven-dried at 80°C for 2 days and recovered in a desiccator with silica gel. Shoot dry weights were measured using the precision balance, and the rosette diameter and leaf blade area were measured using ImageJ version 1.52p.

### 2.3. Growth analysis of plants under in vitro culture on solid media

The shoots and roots of horizontally grown plants were weighed using the precision balance, after which they were oven-dried at 80°C for 2 days and recovered in a desiccator with silica gel. The dry weights of the shoots and roots were measured using the precision balance. The roots of vertically grown plants were scanned at a resolution of 600 dpi to measure the length of primary and lateral roots (>0.3 mm). Root length was measured using ImageJ version 1.52p.

### 2.4. Determination of nutrient elements

Harvested shoots were oven-dried at 80°C for 2 days, and the dried samples were ground using a spoon and recovered in a desiccator with silica gel. For determination of C and N, the dried powder was weighed and encapsulated in a tin boat, and C and N concentrations were measured using a CN analyzer (Vario EL III; Elementar Analysensysteme GmbH, Hanau, Germany). For determination of K, P, S, Ca, Fe, Mg, Mn, Na, B, Co, Cu, Mo, and Zn, the dried samples were weighed and digested using 2 mL of concentrated HNO_3_ followed by the addition of 1 mL of H_2_O_2_ at 120°C until it evaporated. The digested samples were dissolved in 0.08 M HNO_3_ and the element concentrations were measured using inductively coupled plasma–mass spectrometry (Agilent 7800 ICP-MS; Agilent Technologies, Santa Clara, CA, USA) using 2 ppb indium as an internal standard.

### 2.5. Extraction of total RNA

Roots were harvested, immediately frozen using liquid N_2_, and stored at −80°C until use. Frozen samples were ground with a Multi-Beads Shocker (Yasui Kikai, Osaka, Japan) using zirconia beads (diameter: 5 mm). Total RNA was extracted using an RNeasy Plant Mini Kit (Qiagen, Tokyo, Japan) according to the manufacturer’s instructions. For RNA-Seq library preparation, RNA was purified using on-column DNase digestion (Qiagen).

### 2.6. RT-qPCR

Reverse transcription (RT) was performed using ReverTra Ace qPCR RT Master Mix with gDNA Remover (Toyobo, Osaka Japan) according to the manufacturer’s instructions. The synthesized cDNA was diluted 10-fold with distilled water and used in quantitative PCR (qPCR). RT-qPCR was performed on a QuantStudio 1 (Thermo Fisher Scientific) with KOD SYBR qPCR Mix (Toyobo) according to the manufacturer’s instructions. Transcript levels were quantified using a relative standard curve with *ACTIN3* as the internal standard [19]. Primer sequences are shown in Supplementary Table 1.

### 2.7. RNA-Seq analysis

RNA quality was evaluated using a Qubit RNA IQ Assay Kit with a Qubit 4 Fluorometer (Thermo Fisher Scientific, Tokyo, Japan). The RNA samples for which RNA IQs were 9.1–10.0 were used for the following preparation. cDNA libraries were constructed using a NEBNext Ultra II RNA Library Prep with Sample Purification Beads (Code E7775; New England Biolabs, Tokyo, Japan), a NEBNext Poly(A) mRNA Magnetic Isolation Module (Code E7490; New England Biolabs), and a NEBNext Multiplex Oligos for Illumina (Index Primers Set 3) (Code E7710; New England Biolabs) according to the manufacturer’s instructions. These cDNA libraries were sequenced using NextSeq 500 (Illumina, Tokyo, Japan), and the produced bcl files were converted to fastq files using bcl2fastq (Illumina). The resulting sequence data have been stored in the ArrayExpress (https://www.ebi.ac.uk/arrayexpress/) under the accession number (E-MTAB-11843). The reads were analyzed according to the method described by Notaguchi et al. [15] and mapped to the *Arabidopsis* reference (TAIR10) using Bowtie [16] with the following options: “--all --best --strata.” The number of reads mapped to each reference was counted, and the obtained read counts were processed and analyzed using the iDEP ver. 095 integrated web application [17] with the default settings. Pathway enrichment analyses were performed using the Metascape web application with the default settings [18]. Information about read counts is provided in Supplementary Table 2.

### 2.8. Statistical analysis

Unpaired two-tailed Welch’s *t*-tests were conducted using R v.2.15.3.

## 3. Results

### 3.1. Confirmation of root-specific missense mutation of OST2/AHA1

To manipulate the activity of OST2/AHA1 in a root-specific manner, we grafted WT scions onto rootstock originating from WT or *ost2-2D* (Supplementary Fig. 1A). To confirm the missense mutation (G867S) of *OST2/AHA1* reported previously [9], we extracted RNA from the shoots and roots of grafted plants (scion/rootstock: WT/WT and WT/*ost2-2D*). RT-PCR was performed using primers specific to *OST2/AHA1*; after which, the PCR products were sequenced directly. The missense mutation was detected only in the cDNA derived from the WT/*ost2-2D* roots (Supplementary Fig. 1B).

### 3.2. Root-specific activation of OST2/AHA1 enhances shoot growth under nutrient-poor conditions

Grafted plants were transferred to vermiculite pots to analyze the effects of root-specific activation of OST2/AHA1 on shoot growth. Following cultivation for 7 days under nutrient-rich (1/2-MS salts) or nutrient-poor conditions (1/50-MS salts), shoot growth was analyzed. WT/*ost2-2D* plants exhibited increased shoot growth relative to that of WT/WT plants under 1/50-MS but not 1/2-MS conditions (Fig. 1A). Under 1/50-MS conditions, shoot fresh and dry weights were significantly increased by 28% and 31%, respectively, in WT/*ost2-2D* plants than in WT/WT plants (Fig. 1A). Additionally, rosette diameter and leaf blade area were significantly increased by 13% and 29%, respectively, in WT/*ost2-2D* plants than in WT/WT plants under 1/50-MS conditions (Fig. 1A). Under 1/2-MS conditions, however, only the shoot dry weight was significantly increased by 11% in WT/*ost2-2D* plants (Fig. 1A). When the grafted plants were grown in pots containing nutrient-rich soil or in solid media containing 1/2-MS or 1/50-MS, the shoot fresh and dry weights, rosette diameter, and leaf blade area in WT/*ost2-2D* plants were slightly, albeit not always significantly, increased compared with those in WT/WT plants (Supplementary Fig. 2A, B). Relative water content of shoots did not substantially decrease in WT/*ost2-2D* plants compared with those in WT/WT plants regardless of nutritional availability and cultivation style (Supplementary Fig. 3A–C). Thus, root-specific activation of OST2/AHA1 does not cause shoot desiccation due to excessive stomatal opening.

**Fig. 1.**
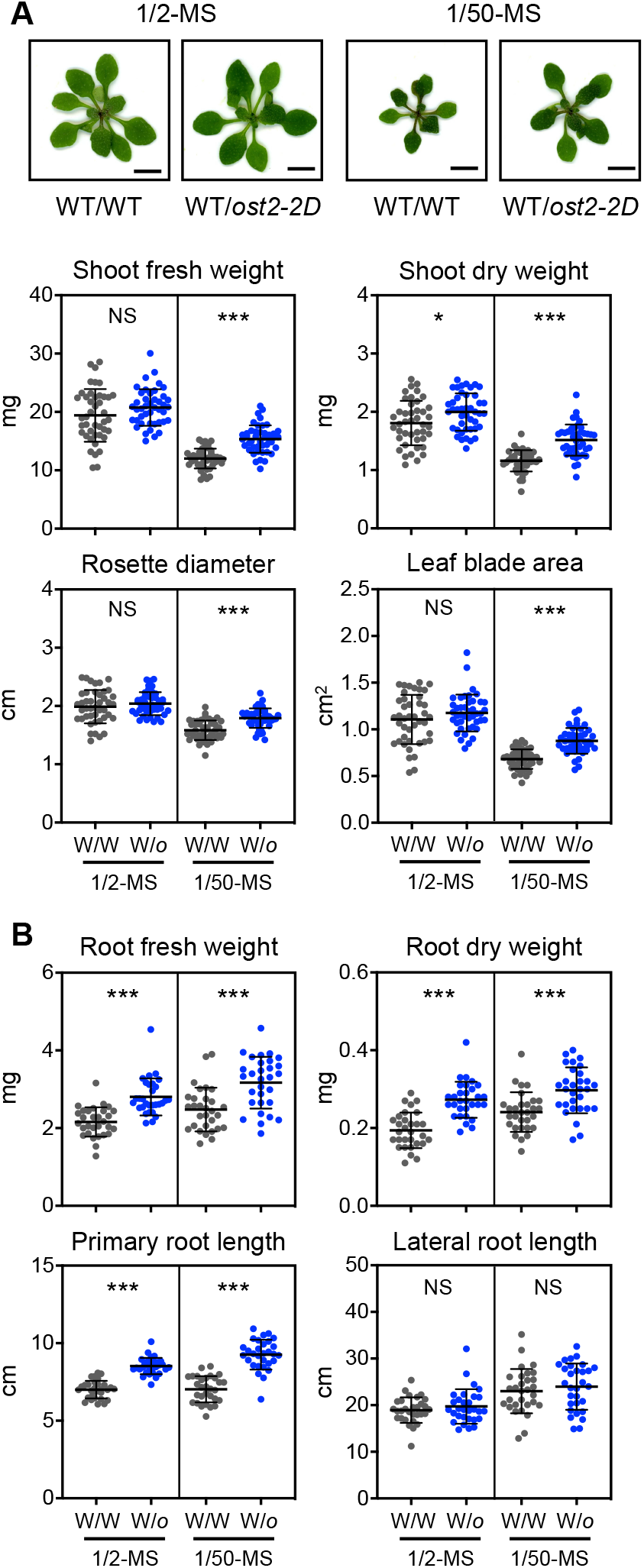
Effects of root-specific activation of OST2/AHA1 on the growth of shoots and roots. (A) Shoot appearance, shoot fresh and dry weights, rosette diameter, and leaf blade area of the grafted plants. Grafted plants were transferred to vermiculite pots and grown for 7 days with a supply of 1/2-MS or 1/50-MS salts. All data pooled from six independent grafting experiments are presented as means ± SD (n = 43–44). (B) Root fresh and dry weights and primary and lateral root lengths of grafted plants. Grafted plants were transferred to 1/2-MS or 1/50-MS media and grown vertically for 3 days. All data pooled from four independent grafting experiments are presented as means ± SD (n = 30). WT/WT (W/W) and *WT/ost2-2D (W/o*) denote plants corresponding to the WT scion grafted on the WT rootstock and the WT scion grafted on the *ost2-2D* rootstock, respectively. \**P* < 0.05; \*\*\**P* < 0.001; NS, not significant at *P* > 0.05 (unpaired two-tailed Welch’s *t*-test). Scale bar: 0.5 cm.

### 3.3. Root-specific activation of OST2/AHA1 enhances root growth irrespective of nutrient availability

Barbez et al. [10] reported that the epidermal cell length of the primary roots of *ost2-2D* seedlings is significantly larger than that of WT plants, suggesting that root growth is altered because of root-specific activation of OST2/AHA1. To confirm this, the root growth of grafted plants grown vertically on solid media containing 1/2-MS or 1/50-MS salts was analyzed. The appearance of the roots in WT/WT plants differed according to nutrient condition, and their fresh and dry weights and lateral root length were generally increased under 1/50-MS conditions than under 1/2-MS conditions (Fig. 1B and Supplementary Fig. 4A). In WT/*ost2-2D* plants relative to in WT/WT plants, root fresh weight and root dry weight were significantly increased by 30% and 41% under 1/2-MS conditions, respectively, and by 28% and 23% under 1/50-MS conditions, respectively (Fig. 1B). The primary root length was significantly increased in WT/*ost2-2D* plants by 22% and 32% under 1/2-MS and 1/50-MS conditions, respectively; however, no significant difference was observed in the lateral root length of the grafted plants (Fig. 1B). In grafted plants grown horizontally on solid media, root fresh and dry weights were increased in WT/*ost2-2D* plants than in WT/WT plants irrespective of nutrient availability (Supplementary Fig. 4B).

### 3.4. Root-specific activation of OST2/AHA1 increases the shoot content of nutrient elements

Zhang et al. [5] reported that constitutive overexpression of *OSA1* significantly increases the content of C, N, K, P, and Ca in the leaves of rice grown in nutrient-rich hydroponic solution. Thus, we determined the content of C and the 14 nutrient elements contained in MS salts in the shoots of grafted plants grown in vermiculite. The content of C, N, K, Ca, S, P, Mg, Na, Mn, B, Co, and Mo was significantly increased in WT/*ost2-2D* shoots than in WT/WT shoots under 1/50-MS conditions (Fig. 2). In particular, the content of macronutrients N, K, Ca, S, and P was consistently increased by 54%, 37%, 24%, 31%, and 44%, respectively, in WT/*ost2-2D* shoots than in WT/WT shoots (Fig. 2). Under 1/2-MS conditions, however, only Na content was significantly increased in the WT/*ost2-2D* shoots than in the WT/WT shoots, although several elements showed increasing trends in the WT/*ost2-2D* shoots (Fig. 2). The shoot concentrations of nutrient elements were generally comparable between the grafted plants regardless of nutrient availability (Supplementary Fig. 5).

**Fig. 2.**
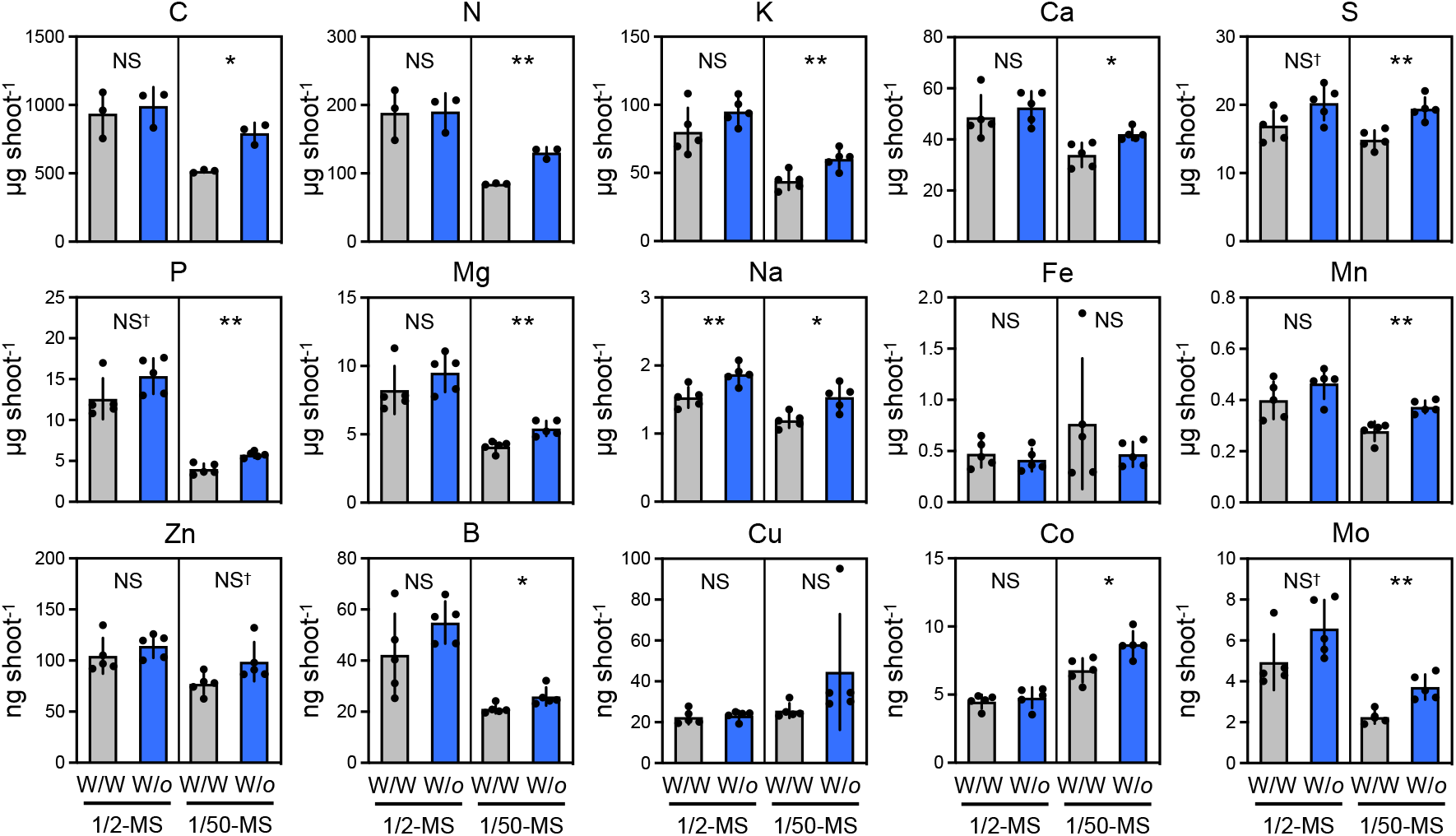
Effects of root-specific activation of OST2/AHA1 on the shoot content of nutrient elements. The shoot content of 15 nutrient elements in the grafted plants, which were transferred to vermiculite pots and grown for 7 days with a supply of 1/2-MS or 1/50-MS salts. Shoot samples harvested from three (for C and N determination) and five (for determination of the other elements) independent grafting experiments were subjected to element analysis. Shoots from 4 to 11 plants in each experiment were pooled as one biological replicate. Data are presented as means ± SD (n = 3 or 5). WT/WT (W/W) and WT/*ost2-2D* (W/*o*) denote the plants corresponding to the WT scion grafted on the WT rootstock and the WT scion grafted on the *ost2-2D* rootstock, respectively. *\*P* < 0.05; *\*\*P* < 0.01; NS^†^ *P* < 0.1; NS, not significant at *P* > 0.05 (unpaired two-tailed Welch’s *t*-test).

### 3.5. Root-specific activation of OST2/AHA1 alters root expression of genes related to biotic responses

To explore the effects of root-specific activation of *OST2/AHA1* on the root transcriptome, we performed RNA-Seq analysis on the roots of grafted plants grown horizontally on solid media containing 1/2-MS or 1/50-MS salts. Hierarchical clustering and principal component analysis (PCA) classified the RNA-Seq data into four groups according to genotype and nutrient condition (Supplementary Fig. 6A, B). The k-means clustering analysis revealed two gene clusters for which the expression was induced (cluster 2) or repressed (cluster 3) in WT/*ost2-2D* roots than in WT/WT roots (Fig. 3A, B and Supplementary Table 2). Each cluster was characterized using pathway enrichment analysis; cluster 2 was enriched in Gene Ontology (GO) terms associated with systemic acquired resistance (SAR) and responses to fungal infection, whereas cluster 3 was enriched in the GO term “lipid X metabolic process” (Fig. 3C). The expression of multiple genes encoding UDP-3-*O*-acyl N-acetylglycosamine deacetylase family proteins LPXC1–5, which contribute to the biosynthesis of lipid X, a precursor of lipid A-like molecules [20], was consistently repressed in WT/*ost2-2D* roots (Supplementary Fig. 7A, B), although the physiological roles played by endogenous lipid A-like molecules are unknown.

**Fig. 3.**
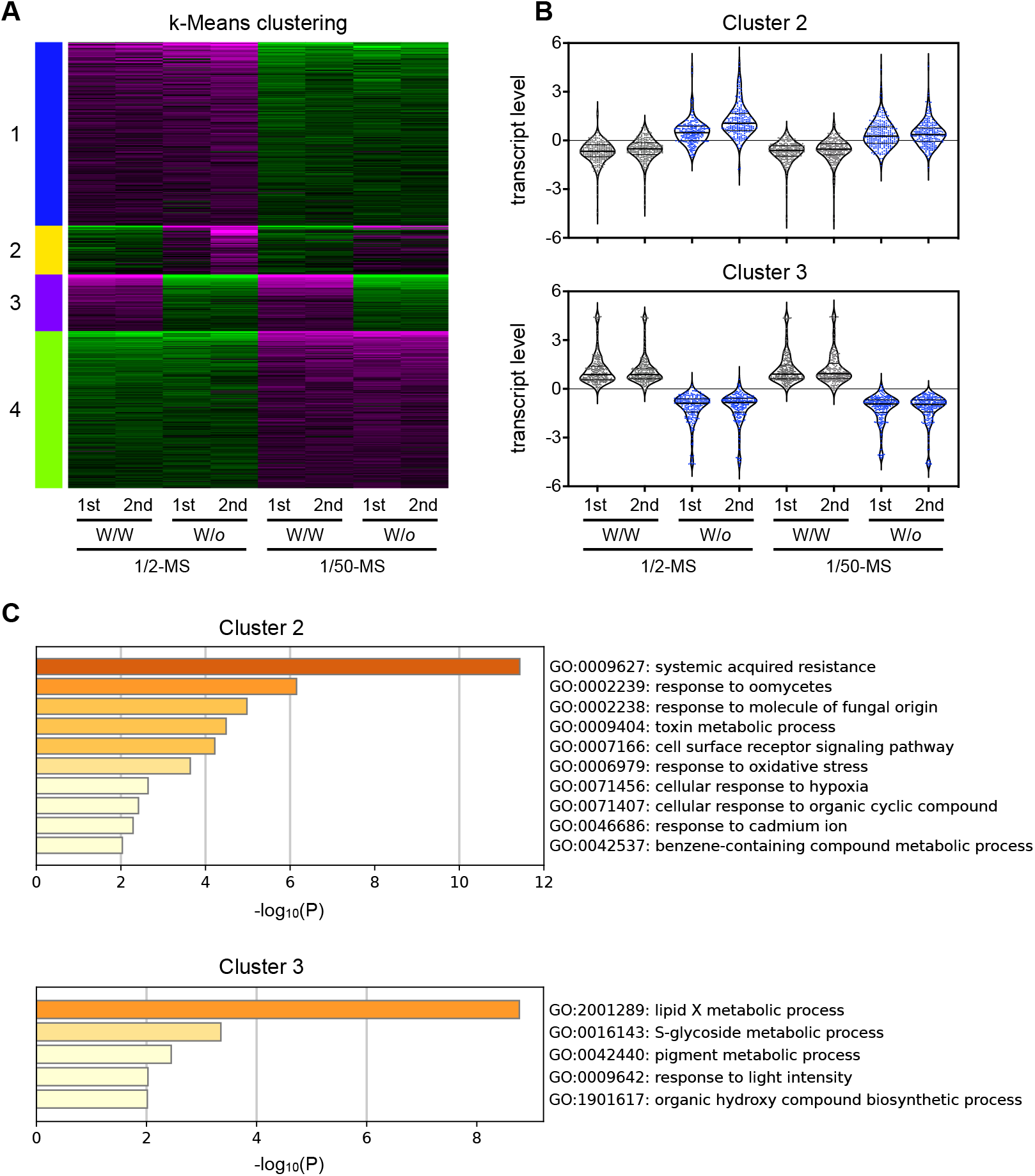
Effects of root-specific activation of OST2/AHA1 on the root transcriptome. (A) Heat map from k-means clustering, (B) violin plots of normalized transcript levels in clusters 2 and 3, and (C) pathway enrichment analyses of the genes in clusters 2 and 3. Grafted plants were transferred to 1/2-MS or 1/50-MS media and grown horizontally for 7 days. Root samples harvested from two independent grafting experiments were subjected to RNA-Seq analysis. Roots from four plants in each experiment were pooled as one biological replicate. Genes for which the standard deviation was ranked in the top 2000 were used for k-means clustering with four clusters, which included 769, 288, 219, and 724 genes (clusters 1–4), respectively. Purple and green represent higher and lower expressions, respectively. Information about genes included in each cluster is provided in Supplementary Table 3. Pathway enrichment analyses were performed using the obtained clusters 2 and 3, and significantly enriched terms are shown. WT/WT (W/W) and WT/*ost2-2D* (W/*o*) denote plants corresponding to the WT scion grafted on the WT rootstock and the WT scion grafted on the *ost2-2D* rootstock, respectively.

## 4. Discussion

The present study demonstrates that root-specific activation of OST2/AHA1 enhances shoot growth and increases the shoot content of all the macronutrient elements and six micronutrient elements in *A. thaliana* cultivated in nutrient-poor vermiculite (Figs. 1A and 2). The root-specific activation of OST2/AHA1 also increased root weight and primary root length in plants grown in solid media (Fig. 1B and Supplementary Fig. 4), corresponding to root cell expansion due to apoplastic acidification, i.e., acid growth [10]. Given that the availability of nutrient elements limits plant vegetative growth under nutrient-poor conditions, the improvement of proton motive force and root growth could facilitate nutrient uptake and shoot growth. In the present study, the concentrations of N, K, P, Mg, B, and Mo in WT/WT shoots were decreased by 19%–53% when nutrient availability was reduced (Supplementary Fig. 5), suggesting that the supply of these elements could limit shoot growth under nutrient-poor conditions. Because the shoot content of N, K, P, Mg, B, and Mo was consistently increased in WT/*ost2-2D* plants than in WT/WT plants (Fig. 2), root-specific activation of OST2/AHA1 may enhance shoot growth by increasing root uptake of these elements. Interestingly, root expression of the high-affinity nitrate uptake transporter *NRT2.1* and iron uptake transporter *IRT1* was higher in WT/*ost2-2D* plants than in WT/WT plants under nutrient-poor conditions (Supplementary Fig. 8A, B). IRT1 has broad specificity for divalent heavy metals other than Fe including Mn, Zn, Co, and Cd under Fe-deficient conditions [21]. Therefore, up-regulation of *NRT2.1* and *IRT1* might have contributed to increases in the shoot content of N, Mn, and Co under nutrient-poor conditions (Fig. 2).

Our transcriptome analysis suggested that root-specific activation of OST2/AHA1 up-regulates genes related to SAR and fungal infection responses in the roots (Fig. 3C). This corresponds to an observation made previously that salicylic acid (SA) levels and expression of SA-responsive marker genes are elevated in the leaves of *ost2-1D* mutants [9], because endogenous SA acts as a signal that mediates local and systemic defense responses to pathogen infection [22]. Interestingly, lowering the pH in external media *per se* is known to induce expression of genes responsive to pathogens, pathogen elicitors, and SA in *A*. *thaliana* roots [23]. Therefore, root-specific activation of OST2/AHA1 may up-regulate expression of these genes through apoplastic acidification [10].

In conclusion, we succeeded in increasing plant vegetative growth under nutrient-poor conditions via root-specific activation of PM H^+^-ATPase using a grafting strategy. Genome editing technologies are applicable for substituting specific DNA sequences for a gene of interest in various plant species; therefore, it may be possible to design and produce rootstocks with activated PM H^+^-ATPase. Overall, our study presents a novel approach to improving NUE in crops and vegetables; thus, it could contribute to the implementation of low-input sustainable agriculture.

## Abbreviations

GO: Gene Ontology
NUE: Nutrient uptake efficiency
PCA: Principal component analysis
PM: plasma membrane
RPM: Reads per million mapped reads
RT: Reverse transcription
SA: Salicylic acid
SAR: Systemic acquired resistance
WT: Wild-type

## Acknowledgments

Seeds of *ost2-2D* were kindly provided by Dr. Jeffrey Leung (CNRS, France). We thank Maki Saiki for ICP-MS analysis. The authors would like to thank Enago (www.enago.jp) for the English language review.

## Funding

This study was supported by a KAKENHI Grant from Japan Society for the Promotion of Science [Grant-in-Aid for Scientific Research (C) No. JP20K05771 to TH] and the Ichimura Foundation for New Technology (to TH).

## Conflict of interest

The authors declare that this study was conducted in the absence of any commercial relationships that could lead to any potential conflicts of interest.

## Notes

### Competing Interest Statement

The authors have declared no competing interest.

## References

[1] S. Savci, An Agricultural Pollutant: Chemical Fertilizer, Int. J. Environ. Sci. Dev. 3 (2012) 77–80. https://doi.org/10.7763/IJESD.2012.V3.191.

[2] V. Kyriakou, I. Garagounis, A. Vourros, et al., An Electrochemical Haber-Bosch Process, Joule 277 (2020) 142–158. https://doi.org/10.1016/j.joule.2019.10.006.

[3] J. Cooper, R. Lombardi, D. Boardman, et al., The future distribution and production of global phosphate rock reserves, Resour. Conserv. Recycl. 57 (2011) 78–86. https://doi.org/10.1016/j.resconrec.2011.09.009.

[4] M. G. Palmgren, Plant plasma membrane H^+^-ATPases: powerhouses for nutrient uptake, Annu. Rev.Plant Biol. 52 (2001) 817–845. https://doi.org/10.1146/annurev.arplant.52.1.817.

[5] M. Zhang, Y. Wang, X. Chen, et al., Plasma membrane H^+^-ATPase overexpression increases rice yield via simultaneous enhancement of nutrient uptake and photosynthesis, Nat. Commun. 12 (2021) 735. https://doi.org/10.1038/s41467-021-20964-4.

[6] M. Hashimoto-Sugimoto, T. Higaki, T. Yaeno, et al., A Munc13-like protein in *Arabidopsis* mediates H^+^-ATPase translocation that is essential for stomatal responses, Nat. Commun. 4 (2013) 2215. https://doi.org/10.1038/ncomms3215.

[7] F. Gévaudant, G. Duby, E. von Stedingk, et al., Expression of a Constitutively Activated Plasma Membrane H^+^-ATPase Alters Plant Development and Increases Salt Tolerance, Plant Physiol. 144 (2007) 1763–1776. https://doi.org/10.1104/pp.107.103762.

[8] J. Falhof, J. T. Pedersen, A. T. Fuglsang, et al., Plasma Membrane H^+^-ATPase Regulation in the Center of Plant Physiology, Mol. Plant 9 (2016) 323–337. https://doi.org/10.1016/j.molp.2015.11.002.

[9] S. Merlot, N. Leonhardt, F. Fenzi, et al., Constitutive activation of a plasma membrane H^+^-ATPase prevents abscisic acid-mediated stomatal closure, EMBO J. 26 (2007) 3216–3226. https://doi.org/10.1038/sj.emboj.7601750.

[10] E. Barbez, K. Dünser, A. Gaidora, et al., Auxin steers root cell expansion via apoplastic pH regulation in *Arabidopsis thaliana*, Proc. Natl. Acad. Sci. USA. 114 (2017) E4884–E4893. https://doi.org/10.1073/pnas.1613499114.

[11] S. D’Alessandro, Y. Mizokami, B. Légeret, et al., The Apocarotenoid β-Cyclocitric Acid Elicits Drought Tolerance in Plants, iScience 19 (2019) 461–473. https://doi.org/10.1016/j.isci.2019.08.003.

[12] J. H. Wong, M. Klejchová, S. A. Snipes, et al., SAUR proteins and PP2C.D phosphatases regulate H^+^-ATPases and K^+^ channels to control stomatal movements, Plant Physiol. 185 (2021) 256–273. https://doi.org/10.1093/plphys/kiaa023.

[13] K. Monden, M. Kojima, Y. Takebayashi, et al., Root-specific Reduction of Cytokinin Perception Enhances Shoot Growth in *Arabidopsis thaliana*, Plant Cell Physiol. (2022) 256–273. https://doi.org/10.1093/pcp/pcac013.

[14] T. Hachiya, T. Oya, K. Monden, et al., A cellophane-supported *Arabidopsis* culture for seamless transfer between different media is useful for studying various nitrogen responses, Soil Sci. Plant Nutr. 67 (2021) 277–282. https://doi.org/10.1080/00380768.2021.1908094.

[15] M. Notaguchi, T. Higashiyama, T. Suzuki, Identification of mRNAs that move over long distances using an RNA-Seq analysis of *Arabidopsis/Nicotiana* benthamiana heterografts, Plant Cell Physiol. 56 (2014) 311–321. https://doi.org/10.1093/pcp/pcu210.

[16] B. Langmead, C. Trapnell, M. Pop, et al., Ultrafast and memory-efficient alignment of short DNA sequences to the human genome, Genome Biol. 10 (2009) R25. https://doi.org/10.1186/gb-2009-10-3-r25.

[17] S. X. Ge, E. W. Son, R. Yao, iDEP: an integrated web application for differential expression and pathway analysis of RNA-Seq data, BMC Bioinform. 19 (2018) 534. https://doi.org/10.1186/s12859-018-2486-6.

[18] Y. Zhou, B. Zhou, L. Pache, et al., Metascape provides a biologist-oriented resource for the analysis of systems-level datasets, Nat. Commun. 10 (2019) 1523. https://doi.org/10.1038/s41467-019-09234-6.

[19] T. Hachiya, J. Inaba, M. Wakazaki, et al., Excessive ammonium assimilation by plastidic glutamine synthetase causes ammonium toxicity in *Arabidopsis thaliana*, Nat. Commun. 12 (2021) 4944. https://doi.org/10.1038/s41467-021-25238-7.

[20] C. Li, Z. Guan, D. Liu, et al., Pathway for lipid A biosynthesis in *Arabidopsis thaliana* resembling that of *Escherichia coli*, Proc. Natl. Acad. Sci. USA. 108 (2011) 11387–11392. https://doi.org/10.1073/pnas.1108840108.

[21] A. Jogawat, B. Yadav, Chhaya, et al., Metal transporters in organelles and their roles in heavy metaltransportation and sequestration mechanisms in plants, Physiol. Plant 173 (2021) 259–275. https://doi.org/10.1111/ppl.13370.

[22] D. F. Klessig, H. W. Choi, D. A. Dempsey, et al., Systemic Acquired Resistance and Salicylic Acid:Past, Present, and Future, Mol. Plant Microbe Interact. 31 (2018) 871–888. https://doi.org/10.1094/MPMI-03-18-0067-CR.

[23] I. D. A. Lager, O. Andréasson, T. L. Dunbar, et al., Changes in external pH rapidly alter plant gene expression and modulate auxin and elicitor responses, Plant Cell Environ. 33 (2010)1513–1528. https://doi.org/10.1111/j.1365-3040.2010.02161.x.

