## Supplemental Figure 1-8 for "Root-specific activation of plasma membrane H^+^-ATPase 1 enhances plant growth and shoot accumulation of nutrient elements under nutrient-poor conditions in *Arabidopsis thaliana*"

### Supplementary Figure 1

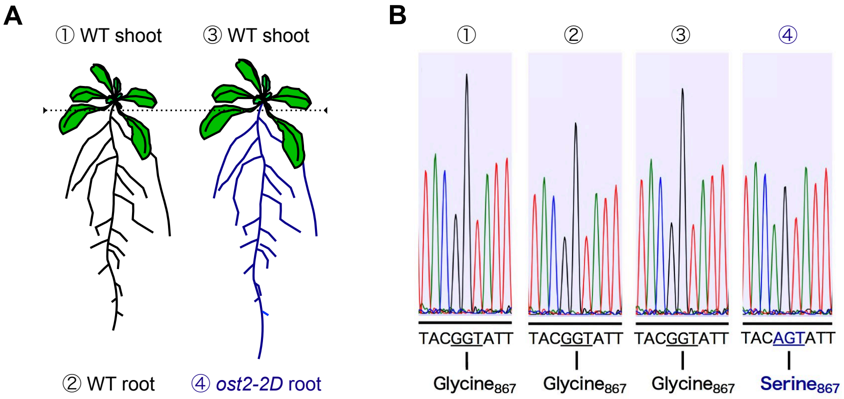


**Supplementary Fig. 1. Root-specific activation of OST2/AHA1 using a micrografting technique.**
(A) Schematic representation of the two types of grafted plants. The dotted line represents the grafted seam in the hypocotyl. (B) Representative results of direct sequencing of the products obtained from RT-PCR using *OST2/AHA1*-specific primers. RNA was extracted from the shoots and roots corresponding to the numbers. The primers used for RT-PCR and direct sequencing are shown in Supplementary Table 1.

### Supplementary Figure 2

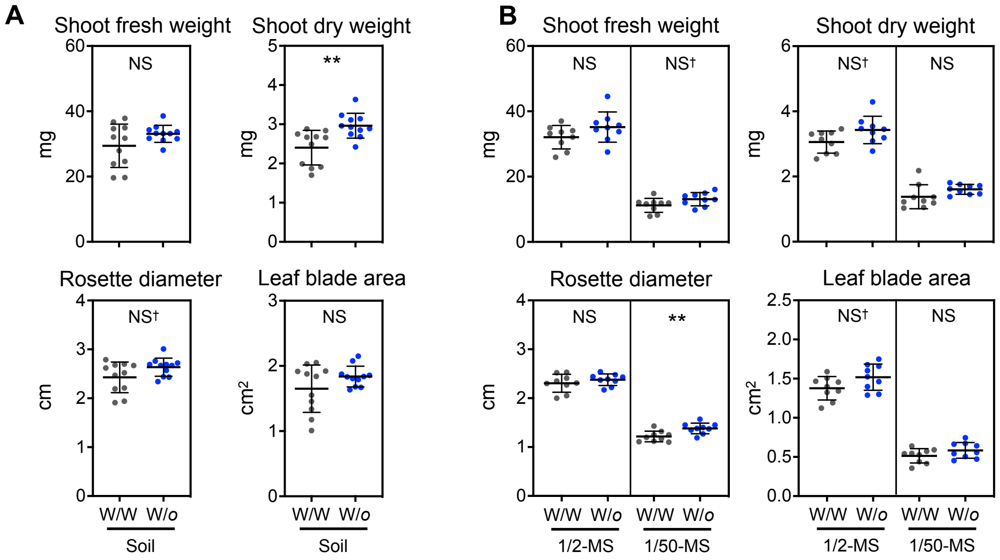


**Supplementary Fig. 2. Effects of root-specific activation of OST2/AHA1 on shoot growth.**

(A) Shoot fresh and dry weights, rosette diameter, and leaf blade area of grafted plants grown under nutrient-rich soil cultivation conditions. Grafted plants were transferred to the soil and grown for 7 days. All data from one grafting experiment are presented as means ± SD (n = 11). (B) Shoot fresh and dry weights, rosette diameter, and leaf blade area of grafted plants grown under in vitro culture conditions. Grafted plants were transferred to 1/2-MS or 1/50-MS media and grown horizontally for 7 days. Data pooled from two independent grafting experiments are presented as means ± SD (n = 9). WT/WT (W/W) and WT/*ost2-2D* (W/*o*) denote plants corresponding to the WT scion grafted on the WT rootstock and the WT scion grafted on the *ost2-2D* rootstock, respectively. $^{**}P < 0.01$; NS† $P < 0.1$; NS, not significant at $P > 0.05$ (unpaired two-tailed Welch's *t*-test).

### Supplementary Figure 3

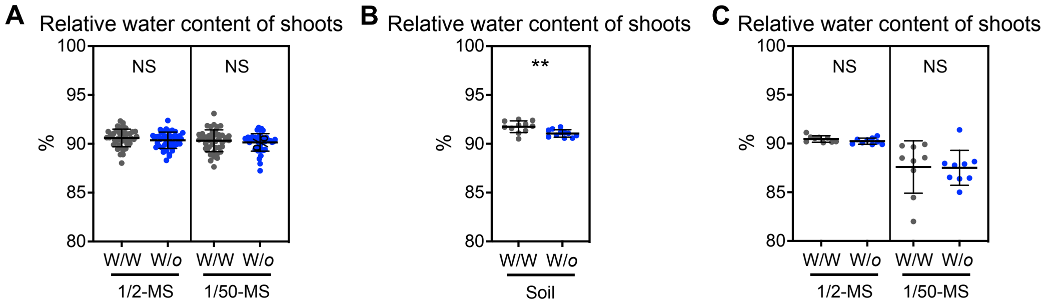


**Supplementary Fig. 3. Effects of root-specific activation of OST2/AHA1 on relative water content of shoots.**
(A) Relative water content of shoots of grafted plants grown under vermiculite cultivation conditions. Grafted plants were transferred to vermiculite pots and grown for 7 days with a supply of 1/2-MS or 1/50-MS salts. Data from six independent grafting experiments are presented as means ± SD (n = 43−44). (B) Relative water content of shoots of grafted plants grown under soil cultivation conditions. Grafted plants were transferred to nutrient-rich soil and grown for 7 days. Data from one grafting experiment are presented as means ± SD (n = 11). (C) Relative water content of shoots of grafted plants grown under i*n vitro* culture conditions. Grafted plants were transferred to 1/2-MS or 1/50-MS media and grown horizontally for 7 days. Data pooled from two independent grafting experiments are presented as means ± SD (n = 9). Relative water contents of shoots (%) were calculated as follows: [(fresh weight − dry weight)/fresh weight] × 100. WT/WT (W/W) and WT/*ost2-2D* (W/*o*) denote plants corresponding to the WT scion grafted on the WT rootstock and the WT scion grafted on the *ost2-2D* rootstock, respectively. ***P* < 0.01; NS, not significant at *P* > 0.05 (unpaired two-tailed Welch's *t*-test).

### Supplementary Figure 4

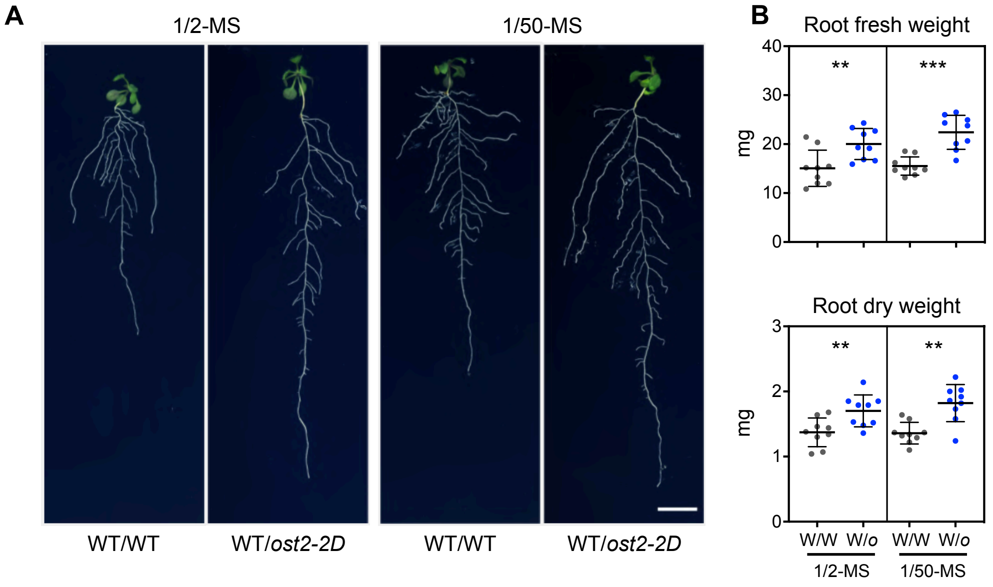


**Supplementary Fig. 4. Effects of root-specific activation of OST2/AHA1 on root growth.**

(A) Representative root appearance in grafted plants grown vertically under in vitro culture conditions. Grafted plants were transferred to 1/2-MS or 1/50-MS media and grown vertically for 3 days. Scale bar: 1 cm. (B) Root fresh and dry weights in grafted plants grown horizontally under in vitro culture conditions. Grafted plants were transferred to 1/2-MS or 1/50-MS media and grown horizontally for 7 days. Data pooled from two independent grafting experiments are presented as means ± SD (n = 9). WT/WT (W/W) and WT/*ost2-2D* (W/*o*) denote plants corresponding to the WT scion grafted on the WT rootstock and the WT scion grafted on the *ost2-2D* rootstock, respectively. $^{**}P < 0.01$; $^{***}P < 0.001$ (unpaired two-tailed Welch's *t*-test).

### Supplementary Figure 5

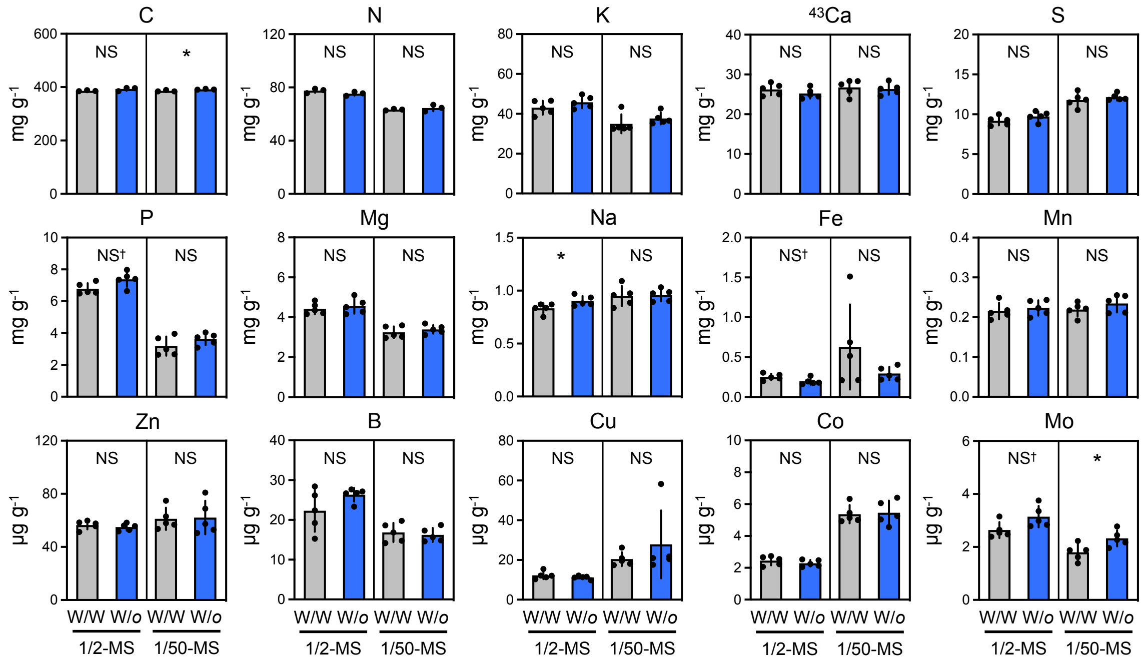


**Supplementary Fig. 5. Effects of root-specific activation of OST2/AHA1 on shoot concentrations of nutrient elements.** The shoot concentrations of 15 nutrient elements in grafted plants, which were transferred to vermiculite pots and grown for 7 days with a supply of 1/2-MS or 1/50-MS salts. Shoot samples harvested from three (for C and N determination) and five (for determination of the other elements) independent grafting experiments were subjected to element analysis. Shoots from 4 to 11 plants in each experiment were pooled as one biological replicate. Data are presented as means ± SD (n = 3 or 5). *$P < 0.05$; NS† $P < 0.1$; NS, not significant at $P > 0.05$ (unpaired two-tailed Welch's *t*-test).

### Supplementary Figure 6

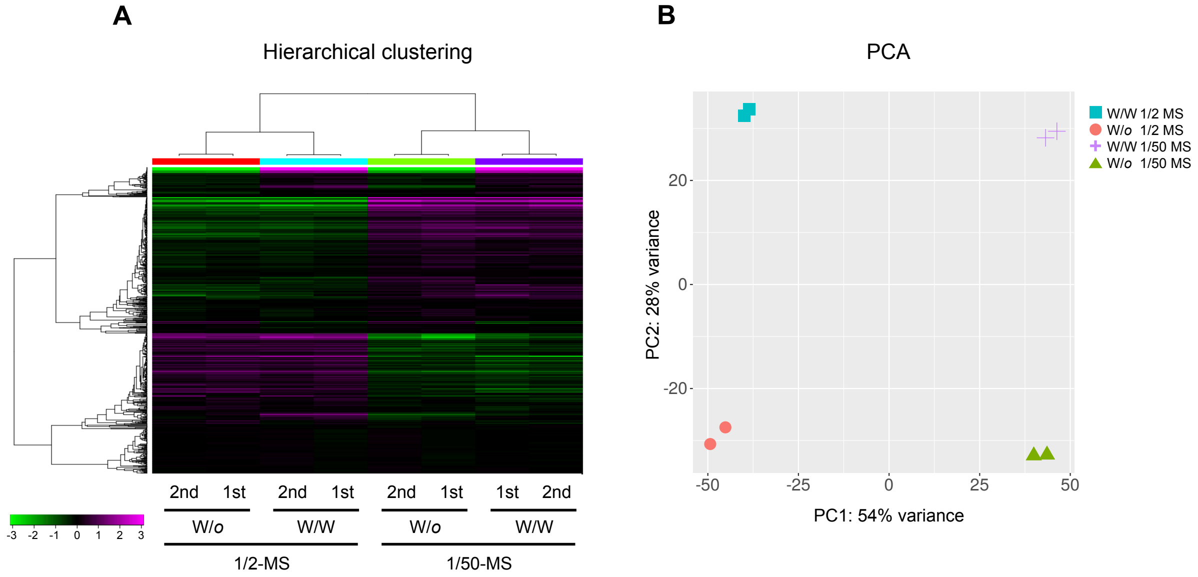


**Supplementary Fig. 6. Effects of root-specific activation of OST2/AHA1 on the root transcriptome.** (A) Heat map from hierarchical clustering and (B) principal component analysis (PCA) results based on RNA-Seq data from grafted plants grown horizontally under in vitro culture conditions. Grafted plants were transferred to 1/2-MS or 1/50-MS media and grown horizontally for 7 days. Root samples harvested from two independent grafting experiments were subjected to RNA-Seq analysis. Roots from four plants in each experiment were pooled as one biological replicate. Hierarchical clustering and PCA were performed using iDEP ver. 095 with the default parameter settings.

### Supplementary Figure 7

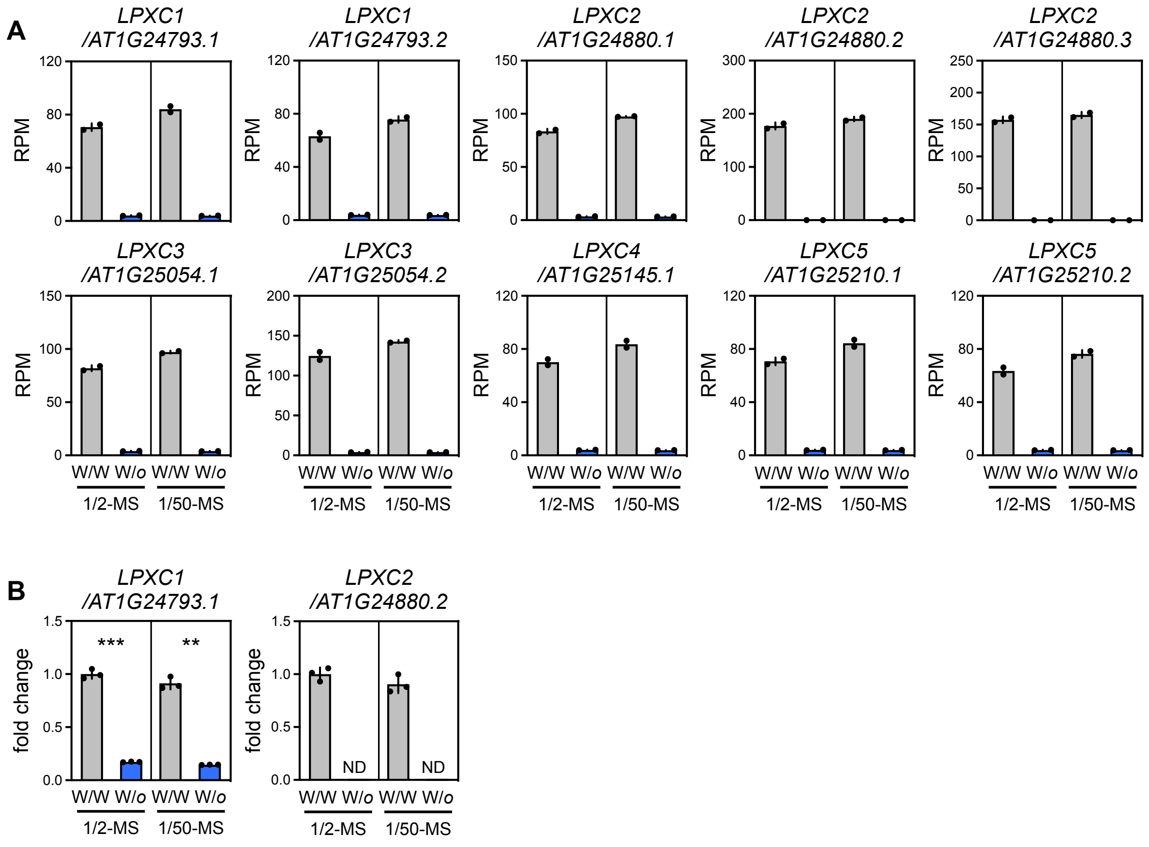


**Supplementary Fig. 7. Effects of root-specific activation of OST2/AHA1 on the root expression of *LPXC* genes.** Relative transcript levels of splice variants of *LPXC* multiple genes were evaluated using (A) RNA-Seq analysis and (B) RT-qPCR. (A) Grafted plants were transferred to 1/2-MS or 1/50-MS media and grown horizontally for 7 days. Root samples harvested from two grafting experiments were subjected to RNA-Seq. Roots from four plants in each experiment were pooled as one biological replicate. Data are presented as means ± SD (n = 2). Transcript levels are shown as reads per million mapped reads (RPM). (B) Grafted plants were transferred to 1/2-MS or 1/50-MS media and grown horizontally for 7 days. Root samples harvested from one grafting experiment were subjected to RT-qPCR. Roots from two plants were pooled as one biological replicate. Data are presented as means ± SD (n = 3). Transcript levels are shown as fold-change values related to the mean transcript level in WT/WT roots under 1/2-MS conditions, given a value of 1. $^{**}P < 0.01$; $^{***}P < 0.001$ (unpaired two-tailed Welch's *t*-test). ND, not detected.

### Supplementary Figure 8

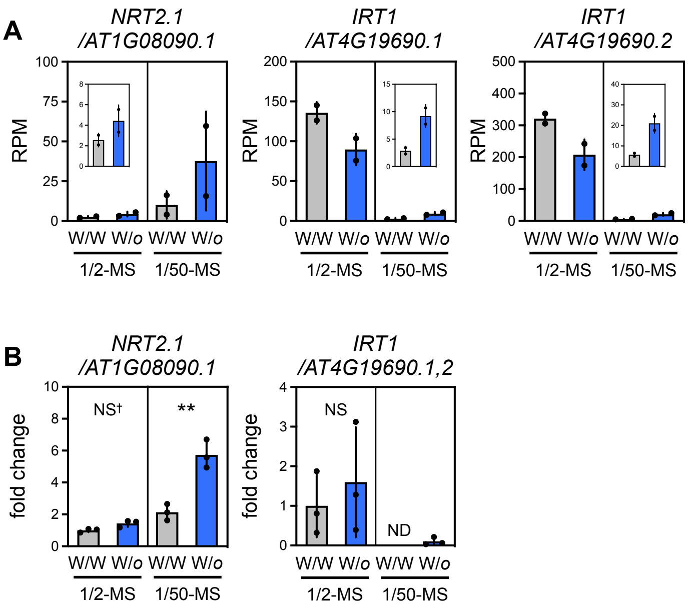


**Supplementary Fig. 8. Effects of root-specific activation of OST2/AHA1 on the root expression of *NRT2.1* and *IRT1*.** Relative transcript levels of splice variants of *NRT2.1* and *IRT1* genes were evaluated using (A) RNA-Seq analysis and (B) RT-qPCR. (A) Grafted plants were transferred to 1/2-MS or 1/50-MS media and grown horizontally for 7 days. Root samples harvested from two grafting experiments were subjected to RNA-Seq. Roots from four plants in each experiment were pooled as one biological replicate. Data are presented as means ± SD (n = 2). Transcript levels are shown as reads per million mapped reads (RPM). (B) Grafted plants were transferred to 1/2-MS or 1/50-MS media and grown horizontally for 7 days. Root samples harvested from one grafting experiment were subjected to RT-qPCR. Roots from two plants were pooled as one biological replicate. Data are presented as means ± SD (n = 3). Transcript levels are shown as fold-change values related to the mean transcript level in WT/WT roots under 1/2-MS conditions, given a value of 1. ***P* < 0.01; NS† *P* < 0.1; NS, not significant at *P* > 0.05 (unpaired two-tailed Welch's *t*-test). ND, not detected.
