## Supplemental Table 1 for "Root-specific activation of plasma membrane H^+^-ATPase 1 enhances plant growth and shoot accumulation of nutrient elements under nutrient-poor conditions in *Arabidopsis thaliana*"

**Supplementary Table 1. Primers used in this study**

| Primer name | Purpose | Primer sequences |
|---|---|---|
| *OST2/AHA1*-RT-F | RT-PCR | GGACAAGTTCGGTGTGAGGT |
| *OST2/AHA1*-RT-R | RT-PCR | GCTATCTCGGCCCTTCTCTT |
| *OST2/AHA1*-S | Direct sequencing | GATCGCAGTGTATGCCGACT |
| *ACTIN3*-qRT-F | RT-qPCR | GGCTAACCGTGAGAAGATGA |
| *ACTIN3*-qRT-R | RT-qPCR | CGACCTGCAAGATCAAGACG |
| *IRT1/AT4G19690.1,2*-qRT-F | RT-qPCR | CCCCGCAAATGATGTTACCTT |
| *IRT1/AT4G19690.1,2*-qRT-R | RT-qPCR | GGTATCGCAAGAGCTGTGCAT |
| *LPXC1/AT1G24793.1*-qRT-F | RT-qPCR | ATCAGACGTTGAATCAATGAGACT |
| *LPXC1/AT1G24793.1*-qRT-R | RT-qPCR | ATCCTGCGAGAGTTTGCTGT |
| *LPXC2/AT1G24880.2*-qRT-F | RT-qPCR | AGAATGCTCCACTCAGCAGC |
| *LPXC2/AT1G24880.2*-qRT-R | RT-qPCR | GCTATTTTTGACAGATGTAGCAGC |
| *NRT2.1/AT1G08090.1*-qRT-F | RT-qPCR | AACAAGGGCTAACGTGGATG |
| *NRT2.1/AT1G08090.1*-qRT-R | RT-qPCR | CTGTGGAAGGAGGCAAGAAC |
